# A novel model of paclitaxel-induced peripheral neuropathy produces a clinically relevant phenotype in mice

**DOI:** 10.1101/2025.02.10.637458

**Authors:** Laura R Osborn, Mary Grace Bishop, Kayleigh A Rodriguez, Dakota M Redling, Elizabeth A Duplechain, Kimberly E Stephens

## Abstract

One of the most common adverse side effects of chemotherapeutics is chemotherapy-induced peripheral neuropathy (CIPN). Paclitaxel, a highly effective chemotherapeutic, is associated with a high incidence of paclitaxel-induced peripheral neuropathy (PIPN) that persists for over a year in 64% of patients and worsens with cumulative PTX dose. Patients experiencing PIPN may reduce the dosage of chemotherapy or halt treatment due to this pain. Current preclinical models have improved our understanding of PIPN but have been ineffective in generating translational therapeutic options. These models administer a single cycle of PTX to induce a PIPN phenotype of mechanical and cold hypersensitivity that resolves within 28 days. However, this does not mirror the clinical dosing regimen or the patient experience of CIPN. In this study, we conduct a comprehensive and longitudinal behavioral profile of our novel model of PIPN in mice where three consecutive cycles of PTX (4 mg/kg, 4 doses/cycle) are given to mimic the clinical administration. Repeated cycles of PTX caused long-lasting mechanical and cold hypersensitivity in male and female C57Bl/6J mice that mirrors clinical observations of persistent CIPN without causing detrimental effects to rodent overall health, normal rodent behavior, or motor function. Our findings support the use of this translational model to facilitate a better understanding of PIPN and the development of effective treatment options. Improved pain management will enable the completion of cancer treatment, decrease health care expenditure, decrease mortality, and improve the quality of life for cancer patients and survivors.

## 1. Introduction

Systemic chemotherapy remains indispensable for the management of many types of cancer, and refinement of chemotherapeutic regimens has improved the survival of cancer patients [18]. While cancer-related mortality rates have fallen, the long-term adverse effects of chemotherapy contribute to a decreased quality of life in cancer survivors. Many chemotherapeutics produce painful peripheral neuropathy that affects up to 96% of patients during treatment and into the cancer survivorship period [9,25]. Affected patients are subjected to an estimated $17,000 of additional healthcare costs, more than 10 additional outpatient visits, and 14% more hospitalizations than cancer patients without chemotherapy-induced peripheral neuropathy (CIPN) [20,29].

Paclitaxel (PTX) is a highly effective taxane used to treat breast, ovarian, and lung cancers. It is administered in 3-6 consecutive cycles over several months with the total dosage and duration of therapy conferring its effectiveness [27,36]. However, PTX is associated with a high incidence of paclitaxel-induced peripheral neuropathy (PIPN) that persists for over a year in 64% of patients and increases in severity with cumulative chemotherapy dose [9,19,21,31,36]. Existing pharmacologic treatments for CIPN are limited in their effectiveness and associated with toxic side effects and additional costs [20,24,29]. Currently, the only effective measures to attenuate CIPN are to reduce the dosage of chemotherapy given or stop treatment, jeopardizing patient survival [9,16].

Rodent PIPN models are pivotal in improving our understanding of PIPN pathogenesis. These models administer a series of four injections of PTX given every other day to mimic a single cycle of PTX with the route of administration, total dosage, and duration of treatment varying among studies [35]. Mechanical and cold hypersensitivity are detectable within three days after the first dose and resolve within four weeks [17,37]. Despite their extensive use, PTX dosing used to generate these preclinical rodent models does not mirror the typical clinical dosing schedule of PTX therapy [13] and the rodent phenotype of rapidly developing and resolving sensitivities is inconsistent with the clinically occurring phenotype of PIPN. Although these models have vastly expanded our understanding of PIPN development, they do not examine PIPN chronification and ultimately have not produced effective CIPN treatments, highlighting the need for improved preclinical CIPN models [40].

We have generated a translational CIPN model in mice where we administer three consecutive cycles of PTX to mimic its clinical administration. In this study, we conduct comprehensive and longitudinal behavioral assessments in mice throughout three consecutive PTX cycles. Similar to patient reports of PIPN, where dosage and length of therapy increase the severity of painful neuropathy [9,42], mice experienced an increased duration of mechanical and cold hypersensitivity after repeated cycles of PTX compared to earlier cycles. Our translational PIPN model will improve our understanding of PIPN development, resolution, and chronification by allowing us to mimic the clinically used dosing regimen and produce a clinically relevant phenotype in rodents that will facilitate the development of novel effective treatment options.

## 2. Methods

### 2.1 Animals

Adult male and female C57BL/6J mice (8 weeks) were purchased from Jackson Laboratories (Bar Harbor, ME) and housed in a room with controlled temperature (22 ± 1 °C) and humidity (60 ± 10%) with a 12-hour light/dark cycle. Standard rodent chow and water were available *ad libitum*. Mice were allowed to acclimate to the housing facility for a minimum of 48 hours prior to any experimental procedures. Each mouse was randomly assigned to a treatment group upon arrival and all mice in a cage (4-5/cage) received the same experimental treatment. All exp’erimental procedures were performed during the animal’s light cycle (7:00 am to 7:00 pm). Experimenters performing behavioral testing were blinded to group assignment. All procedures involving animals were reviewed and approved by the UAMS Institutional Animal Care and Use Committee (IACUC) and were performed in accordance with the National Institute of Health’s Guide for the Care and Use of Laboratory Animals.

### 2.2 Paclitaxel

Paclitaxel (Athenex, Schaumburg, IL) was obtained from Arkansas Children’s Hospital Pharmacy as 6 mg/kg paclitaxel dissolved in 527 mg Polyoxyl 35 Castor Oil NF, 49.7% (V/V) United States Pharmacopeia Grade Dehydrated Alcohol and 2 mg Citric Acid. Immediately prior to administration, paclitaxel was diluted to 0.5 mg/mL in sterile 0.9% sodium chloride. Control mice received vehicle only [5% ethanol, 5% Cremophor EL (Merck KGaA, Darmstadt, Germany), 95% sterile 0.9% sodium chloride]. One cycle consisted of four intraperitoneal (IP) paclitaxel injections (4mg/kg) given every other day for a cumulative dosage of 16 mg/kg [14,37]. Each animal received a total of three cycles (cumulative dosage of 48 mg/kg).

### 2.3 Experimental design

Mice were randomly assigned to PTX or vehicle groups. Female mice were randomly assigned to one of two cohorts: a behavioral cohort (n=16) or an open field cohort (n=64). Male mice were assigned to the behavioral cohort (n=16). As shown in Figure 1, the first cycle of PTX or vehicle was administered to all mice following baseline assessments of the behavioral cohort. Once mechanical sensitivity returned to baseline in the behavioral cohort, a second cycle of PTX or vehicle was administered to all mice (i.e., 28-30 days after the first PTX dose of cycle 1). A third cycle was administered once mechanical sensitivity had returned to baseline in the behavioral cohort (i.e., 28-30 days after the first PTX dose of cycle 2). Mice in the behavioral cohort underwent comprehensive and longitudinal assessments for mechanical and cold sensitivity and rotarod and burrowing behavior for the duration of the study. They were also weighed weekly, and their food intake was monitored throughout the study. Mice in the open field cohort were assessed for open field behavior during mechanical hypersensitivity for each cycle (n=8/group/time point). Male mice in the behavioral cohort were assessed for open field behavior during mechanical hypersensitivity during cycle 3 (n=8/group). Mechanical sensitivity was assessed prior to the open field test. These mice were euthanized following completion of the open field test and blood was used for hematological analysis as detailed below.

**Figure 1.**
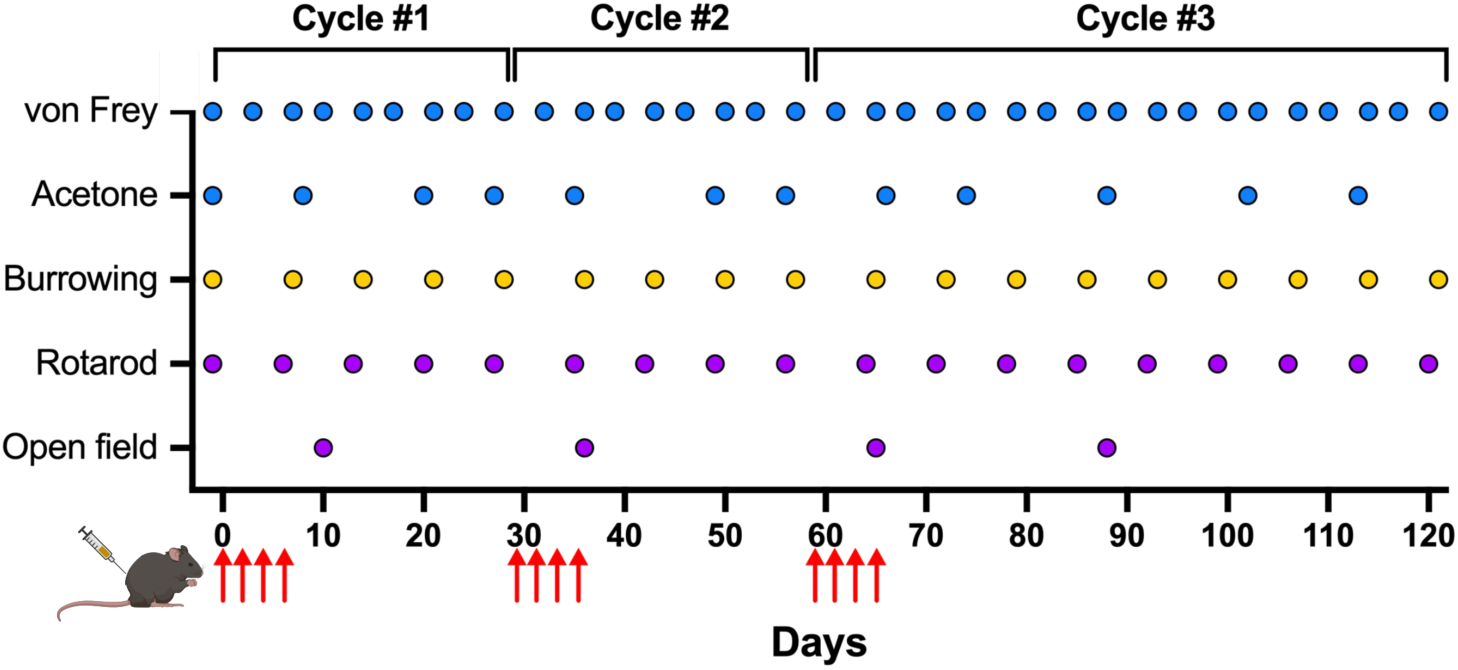
Schematic of experimental design. Each mouse received 3 consecutive cycles of PTX (4 mg/kg/dose) or vehicle. Each cycle consisted of 4 doses (red arrows) and subsequent cycles were given once mechanical sensitivity for all animals returned to baseline. Behavioral assessments of evoked responses (blue), normal rodent behavior (yellow), and spontaneous behavior (purple) were conducted throughout PTX administration.

### 2.4 Behavioral assessments

#### 2.4.1 Mechanical

Mechanical sensitivity was assessed using von Frey monofilaments (Stoeling Co., Wood Dale, IL) following the low-high assessment method [7] at baseline and twice weekly for 16 weeks. Prior to baseline assessment, mice were habituated for >1 hour/day for 2 days to the testing apparatus which consisted of individual plexiglass boxes on a wire mesh floor. On days of testing, mice were habituated to the apparatus for >30 minutes before testing. Paw withdrawal frequency was assessed to low-force (0.6 gram) and high-force (1.4 gram) mechanical stimulation using von Frey monofilaments. The designated monofilament was applied to the plantar surface of the hind paw for 1 second. The stimulation was repeated for 10 trials on each hind paw for both filaments with >5 minutes between each stimulation of an individual paw. Each trial was scored to indicate a positive or negative response by the mouse. A positive response was defined as the abrupt lifting of the paw with shaking, extended lifting, and/or licking of the foot. The percent paw withdrawal frequency (%PWF) was calculated as 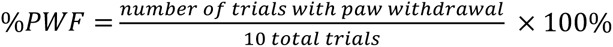. At each time point, %PWF for each animal was determined by averaging the %PWF of the right and left hind paws.

### 2.4.2 Cold

Cold sensitivity was assessed using the acetone test [5,41] at baseline, twice weekly for 16 weeks, and then once weekly for four weeks. Mice were placed in individual plexiglass boxes on a wire mesh floor and habituated for >30 minutes prior to each assessment. Acetone (∼50 uL) was applied to the hind paw via a syringe and time spent reacting (TSR) to the stimulation (e.g., licking, lifting, and/or shaking) was recorded for the 60 seconds immediately following stimulation. Stimulation with acetone was repeated for 5 trials on each hind paw with >5 minutes between each stimulation of an individual paw. At each time point, TSR (s) for each animal was determined by averaging the TSR of the right and left hind paws for five trials.

#### 2.4.3 Thermal

Thermal sensitivity was assessed using the Hargreaves test [15] at baseline and once weekly for 4 weeks. Mice were placed in individual plexiglass boxes on a glass floor and habituated for >30 minutes prior to each assessment. Thermal stimulation from an infrared source was focused below the hind paw and paw withdrawal latency (PWL) to the stimulation (licking, lifting, and/or shaking) was recorded automatically. To avoid damage to the hind paw, the maximum PWL was 20 seconds. Stimulation was repeated for 3 trials on each hind paw with >5 minutes between each stimulation of an individual paw. PWL (s) for each animal was determined by averaging the PWL of the right and left hind paws for three trials.

#### 2.4.4 Burrowing

Pain-depressed burrowing performance was assessed using the burrowing test [10,33] at baseline and once weekly for 16 weeks and was conducted in the same room where the mice were housed. Polyvinyl chloride (PVC) tubes (5.1 cm in diameter x 12.7 cm in length) were filled with 80 grams of corncob bedding. The open end of the burrow was elevated 5 cm from the floor of the cage by 2 metal machine screw legs to prevent spilling of the burrow contents. Prior to testing, mice were habituated to burrows for 12 hours in their home cage which allowed cage mates to learn burrowing behavior through social facilitation. All cages exhibited burrowing behavior and were included in testing. Following habituation, mice underwent baseline testing to confirm all individual mice displayed burrowing behavior which was defined as the ability to burrow >50% of the burrow contents in the testing time. All mice displayed burrowing behavior and were included in subsequent testing. In the testing phase, mice were placed individually in a clean cage with the burrow for 20 minutes. At the conclusion of the test, the amount of bedding burrowed (grams) for each mouse was determined after weighing the remaining bedding. Each burrowing test was filmed to determine each mouse’s latency to burrow.

#### 2.4.5 Proprioception

Motor function was measured using the rotarod test [11,34] at baseline and once weekly for 16 weeks. Mice were acclimated to the testing room for >15 minutes prior to testing. Mice were first trained on the rotarod (Rotamex-5, Columbus Instruments, Columbus, OH) for three consecutive days to ensure the ability to walk forward on the rotating rod. During training, mice were required to remain on the rod for 1 minute at 4 rpm. By the third day of training, all mice were able to remain on the rod, therefore no mice were excluded from subsequent testing. In the testing phase, the mice were placed on the rod rotating at 4 rpm for 10 seconds. The rotation speed then ramped to 40 rpm over 5 minutes (acceleration of 0.6 rpm every 5 seconds). The latency for the mouse to fall from the rod during the acceleration period was automatically recorded. Each mouse was tested three times with >15 minutes between trials. For each animal, the latency to fall and rotation speed at fall were averaged for all three trials. If the animal fell off the rod at 4 rpm, before ramping began, they were returned to the rod. After a second fall during this period the trial would end, and their latency to fall was recorded as 0 seconds (4 rpm at fall).

#### 2.4.6 Open field

Movement was assessed with the open field test [30,32] during mechanical hypersensitivity (days 7-10 following the first dose of PTX) for each cycle as well as 30 days following the first dose of the third cycle of PTX. Without prior habituation, each mouse was positioned at the center of the open field apparatus (40 cm x 40 cm x 40 cm) and allowed to move freely for 10 minutes. The movements of the mouse were continuously captured by a video camera positioned above the apparatus. The video files were analyzed using Any-Maze (Stoeling Co., Wood Dale, IL) for the total distance traveled, average speed, time spent moving, and time in the center, corner, and perimeter areas. Following each trial, the apparatus was thoroughly cleaned with 70% ethanol before the next mouse was tested. Each mouse was tested once, and testing was conducted under red light.

### 2.5 Blood analysis

Trunk blood was collected during the period of peak mechanical hypersensitivity (days 7-10 following the first dose of PTX) for each cycle, 30 days following the first dose of the third cycle of PTX, and following the resolution of mechanical hypersensitivity after each PTX cycle. Blood was analyzed using the DxH 500 Series Hematology Analyzer (Beckman Coulter, Brea, CA) and values were compared to Jackson Laboratory Physiological Data Summary for C57BL/6J female mice [1].

### 2.6 Statistical analysis

All data are represented as mean ± SD. The sample size for all experiments is noted in the figure legends. Statistical analysis was performed using GraphPad Prism version 10.4.1 for Windows, GraphPad Software, Boston, Massachusetts USA, www.graphpad.com. Data were analyzed by 2-way ANOVA followed by Tukey’s post hoc test for multiple comparisons or by Student’s t test for single comparisons. A p-value <0.05 was considered statistically significant.

## 3. Results

### 3.1 Effects of repeated PTX cycles on rodent health

Our novel model of PIPN utilizes three consecutive cycles of PTX during which we conducted longitudinal and comprehensive behavioral assessments (Fig. 1). We monitored animal overall well-being throughout the study. As expected, body weight significantly increased over time for both PTX and vehicle groups of male [*F*(4.287, 60.02) = 99.23, p<0.0001] and female [*F*(3.964, 55.50) = 39.53, p<0.0001] mice (Fig. 2A). While we found no difference in body weight between PTX and vehicle female mice, PTX did reduce weight gain in male mice compared to vehicle [*F*(1, 14) = 23.31, p=0.0003] (Fig. 2A). There were no apparent differences in food intake between PTX and vehicle groups for male or female mice (Fig. 2B), suggesting that PTX did not induce nausea or loss of appetite. To further examine physiological changes that may occur due to repeated PTX cycles in our model, we performed hematological analysis on blood obtained from female mice during mechanical hypersensitivity and at resolution following each cycle of PTX. We found no differences in red blood cell (RBC), white blood cell (WBC), and platelet counts, mean corpuscular volume, hemoglobin or mean corpuscular hemoglobin, or the percent contents of lymphocytes, monocytes, neutrophils, eosinophils, or basophils between the PTX and vehicle groups (Supp Fig 1). However, average values of RBC, WBC, hemoglobin, and platelets for both PTX and vehicle female mice were lower than the normal range.

**Figure 2.**
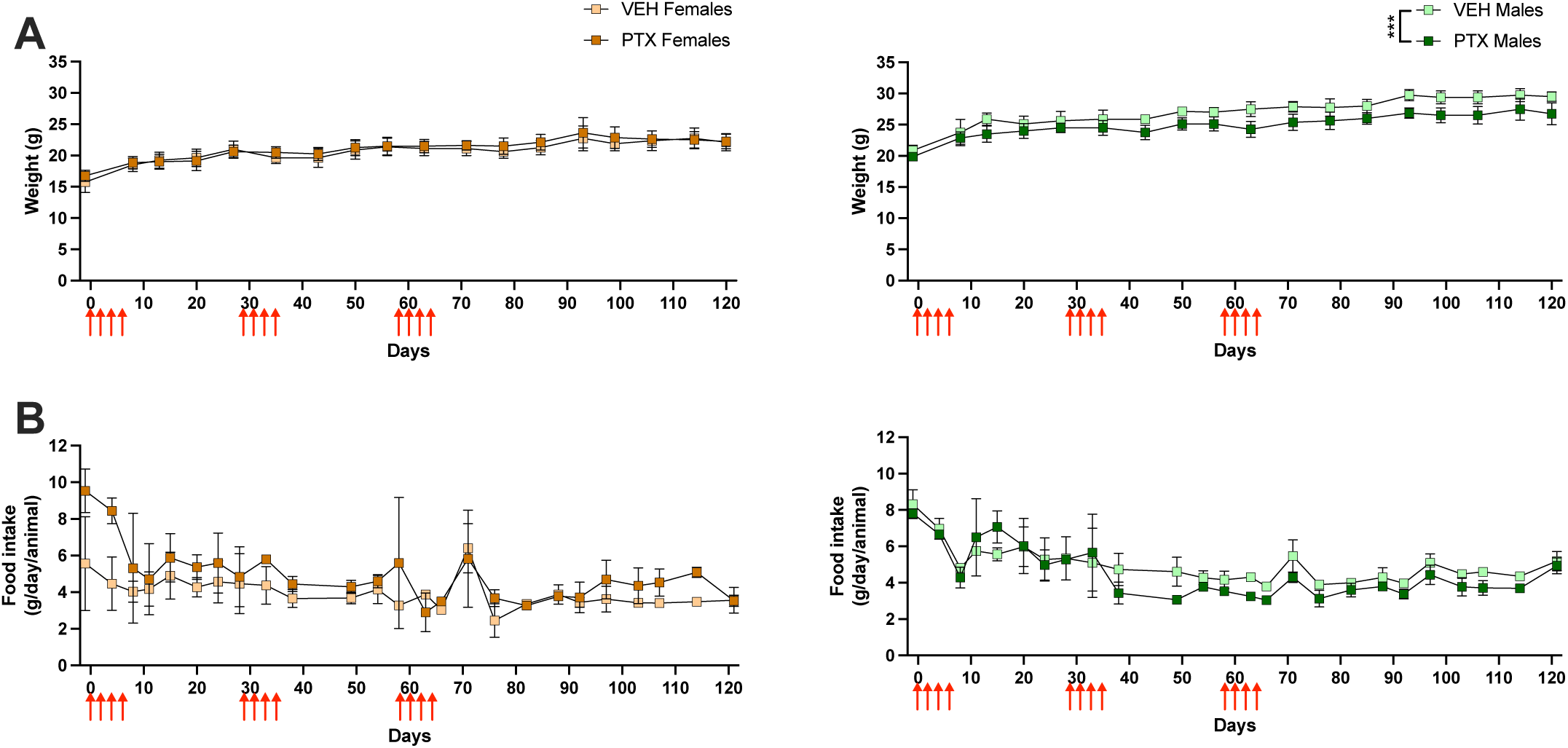
General well-being of mice throughout study. **A)** Weight (grams) throughout administration of 3 PTX cycles (red arrows) in female (left) and male (right) mice (n=8/group). **B)** Food intake (grams/day/animal) throughout administration of 3 PTX cycles in female (left) and male (right) mice (n=2 cages/group). Data is represented as mean±SD. *** p<0.001

### 3.2 Repeated PTX cycles do not alter burrowing behavior

The burrowing test was used to monitor normal rodent behavior throughout the administration of PTX. We found no difference in the amount of bedding burrowed between PTX and vehicle groups for male or female mice throughout the study. However, the total amount of bedding burrowed during each assessment increased for male [*F*(3.179, 44.51) = 5.898, p=0.0015] and female [*F*(4.512, 62.90) = 8.186, p<0.001] mice, as the test was repeated over time (Fig. 3A). Similarly, the latency to begin burrowing was unaffected by PTX in male or female mice and we found a significant decrease in the latency to begin burrowing over time for males [*F*(4.471, 62.59) = 6.700, p<0.0001] and females [*F*(6.105, 82.24) = 4.355, p=0.0007] (Fig. 3B).

**Figure 3.**
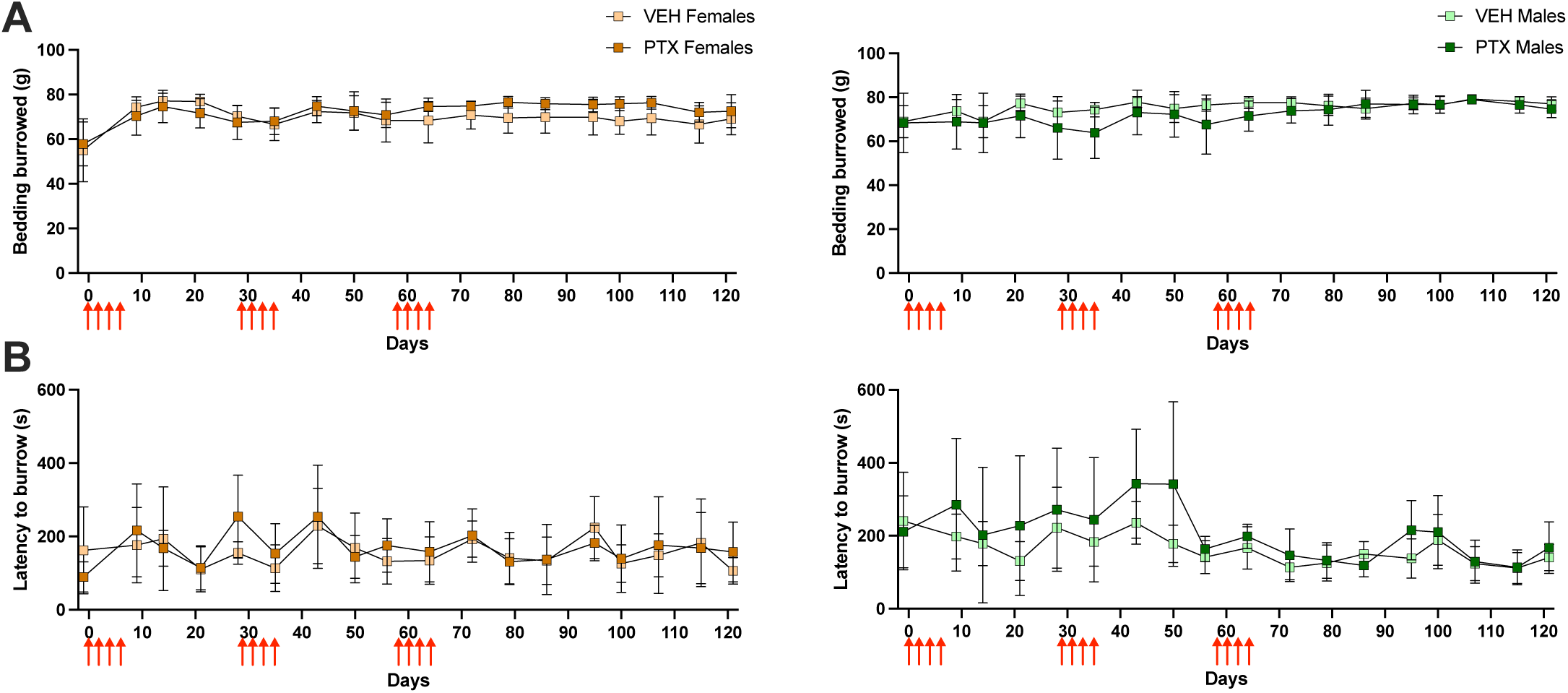
Burrowing behavior after multiple PTX cycles. **A)** Amount of bedding burrowed (grams) and **B)** latency to begin burrowing (seconds) throughout administration of 3 PTX cycles (red arrows) in female (left) and male (right) mice. (n=8/group). Data is represented as mean±SD.

### 3.3 Repeated PTX cycles increase the duration of mechanical hypersensitivity

We used the von Frey test to examine how mechanical sensitivity is altered due to repeated PTX cycles. Mechanical sensitivity was significantly increased in the PTX group following each cycle of PTX in female mice to low-force [*F*(1, 14) = 85.39, p<0.0001] and high-force [*F*(1, 14) = 87.18, p<0.0001] filaments (Figs. 4A-B). Male mice also developed mechanical hypersensitivity to low-force [*F*(1, 14) = 34.39, p<0.0001] and high-force [*F*(1, 14) = 106.6, p<0.0001] filaments following each cycle of PTX (Figs. 4A-B). This hypersensitivity peaked 7-10 days following the first dose of PTX in each cycle and resolved within 4 weeks following the first PTX dose in the first two cycles. However, mechanical hypersensitivity didn’t resolve until 8 weeks following the third PTX cycle (Figs. 4A-B).

**Figure 4.**
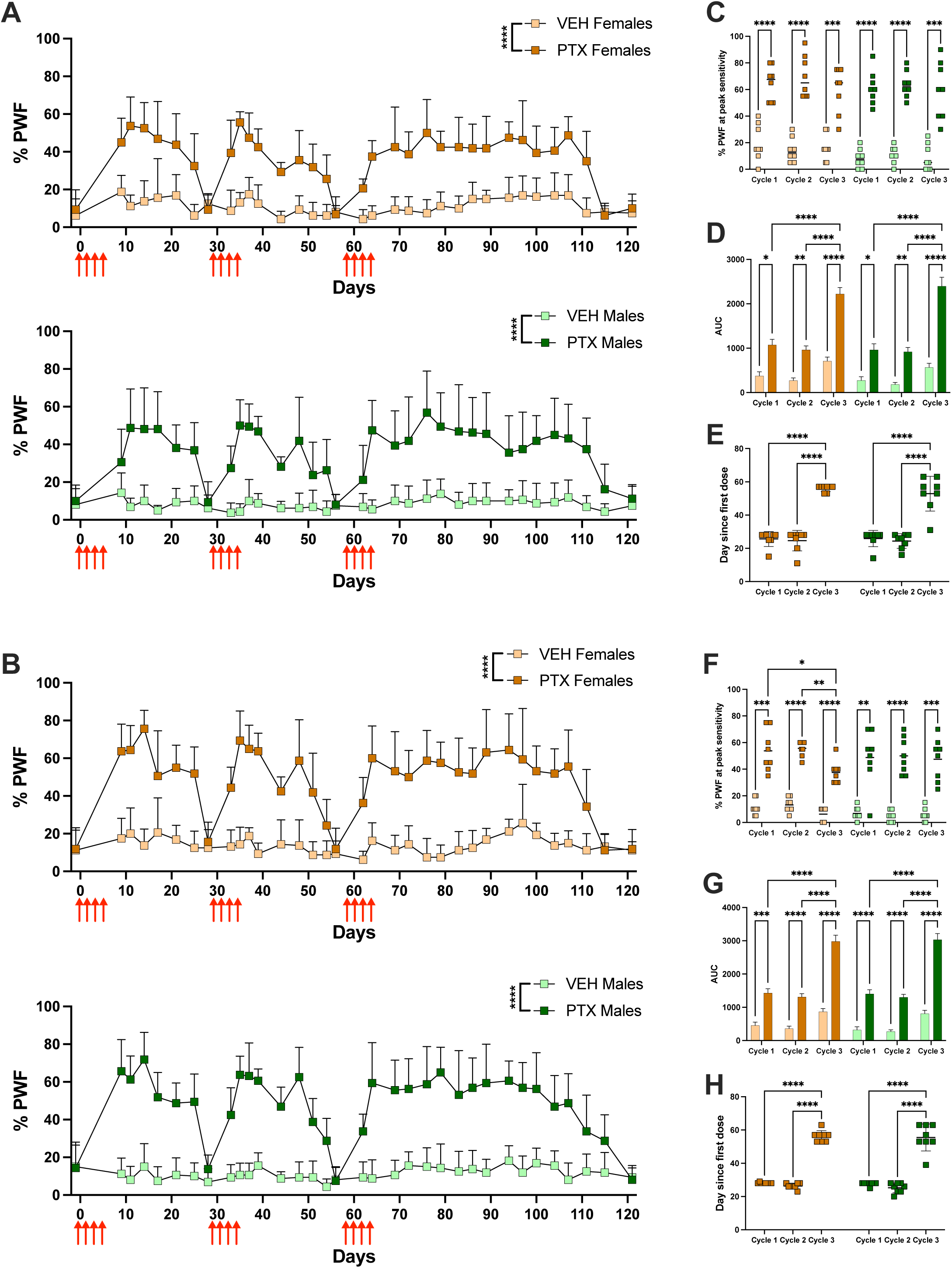
Increased duration of mechanical hypersensitivity after multiple PTX cycles. %PWF to **A)** low (0.6 g) and **B)** high (1.4 g) force filaments throughout the administration of 3 PTX cycles (red arrows) in female (top) and male (bottom) mice. Height of %PWF to **C)** low (0.6 g) and **D)** high (1.4 g) force filaments during days 7-10 following the administration of each PTX cycle in female and male mice. Area under the curve (AUC) of mechanical sensitivity to **E)** low (0.6 g) and **F)** high (1.4 g) force filaments following the administration of each PTX cycle in female and male mice. Duration of mechanical sensitivity (days) to **G)** low (0.6 g) and **H)** high (1.4 g) force filaments following the administration of each PTX cycle in female and male mice. (n=8/group). Data is represented as mean±SD. * p < 0.05, ** p < 0.01, *** p<0.001, and **** p < 0.0001.

The intensity of mechanical hypersensitivity, which we define as the %PWF during peak hypersensitivity (days 7-10) following each PTX cycle, was significantly higher in the PTX group compared to the vehicle group [*F*(3, 28) = 63.48, p<0.0001] to the low-force filament. The intensity of mechanical hypersensitivity was lower in female mice following the third PTX cycle [*F*(1.356, 18.98) = 4.969, p=0.0288] compared to cycles 1 (p=0.0148) and 2 (p=0.0027) but was not different in male mice among the three PTX cycles (Fig. 4C). PTX mice had a significantly higher intensity of mechanical hypersensitivity to the high-force filament [*F*(3, 28) = 80.79, p<0.0001] compared to vehicle mice, and that did not differ between the three cycles (Fig. 4D). There was no effect of sex on the intensity of mechanical hypersensitivity to the low or high-force filaments (Fig. 4C, D), indicating male and female mice experienced similar degrees of mechanical hypersensitivity.

The area under the curve (AUC) of %PWF from stimulation by the low and high-force filaments was significantly increased in the PTX mice compared to the vehicle [*F*(2, 996) = 53.66, p<0.0001 and *F*(2, 996) = 89.58, p<0.001] (Fig. 4E, F). Mechanical hypersensitivity during the third PTX cycle for male and female mice had a greater AUC for stimulation by the low-force [*F*(2, 996) = 62.13, p<0.0001] and high-force [*F*(2, 996) = 88.92, p<0.0001] filaments compared to cycles 1 (p<0.0001) and 2 (p<0.0001) (Fig. 4E, F). There were no differences in the AUC for the mechanical hypersensitivity between male and female mice (Fig. 4E, F). Following the third cycle of PTX, we found that male and female mice had a significantly increased duration of mechanical hypersensitivity to the low-force [*F*(1.979, 27.71) = 240.3, p<0.0001] and high-force [*F*(1.979, 27.71) = 240.3, p<0.001] filaments compared to cycles 1 (p<0.0001) and 2 (p<0.0001) (Fig. 4G, H). However, we found no differences in the duration of PTX-induced mechanical hypersensitivity between male and female mice (Fig. 4G, H).

### 3.4 Repeated PTX cycles increase the duration of cold hypersensitivity

We utilized the acetone test to examine the development of cold hypersensitivity following three consecutive cycles of PTX. Cold hypersensitivity was significantly increased following PTX in female [*F*(1, 14) = 52.14, p<0.0001] and male [*F*(1,14) = 60.00, p<0.0001] mice compared to vehicle mice (Fig. 5A). Cold hypersensitivity developed within 7-10 days following PTX, but did not return to baseline within 4 weeks (Fig. 5A). Cold hypersensitivity developed within 7-10 days following PTX and resolved after 6 weeks in female mice that received a single cycle of PTX (data not shown). Since cold hypersensitivity did not have consistent peaks between cycles, the TSR for all the time points within a cycle were averaged and compared. The average TSR was significantly higher in PTX mice compared to the vehicle for both males and female mice [*F*(3, 28) = 41.02, p<0.0001] (Fig. 5B). The average TSR was significantly increased during cycles 2 and 3 compared to cycle 1 [*F*(2, 56) = 22.18, p<0.0001] for both male and female mice (Fig. 5B). Interestingly, PTX increased the average TSR for females more than males during cycle 1 (p=0.0041), cycle 2 (p<0.0001), and cycle 3 (p=0.0019), indicating that female mice exhibited greater cold hypersensitivity in response to PTX than males. The AUC for TSR from acetone stimulation was significantly increased in PTX mice compared to vehicle [*F*(3, 380) = 17.19, p<0.0001] (Fig. 5C). Cold hypersensitivity during the third PTX cycle for both male and female mice had a greater AUC [*F*(2, 380) = 114.5, p<0.0001] compared to cycles 1 (p<0.0001) and 2 (p<0.0001) (Fig. 5C). Thermal hypersensitivity did not develop in female mice following a single cycle of PTX (Supp Fig. 2).

**Figure 5.**
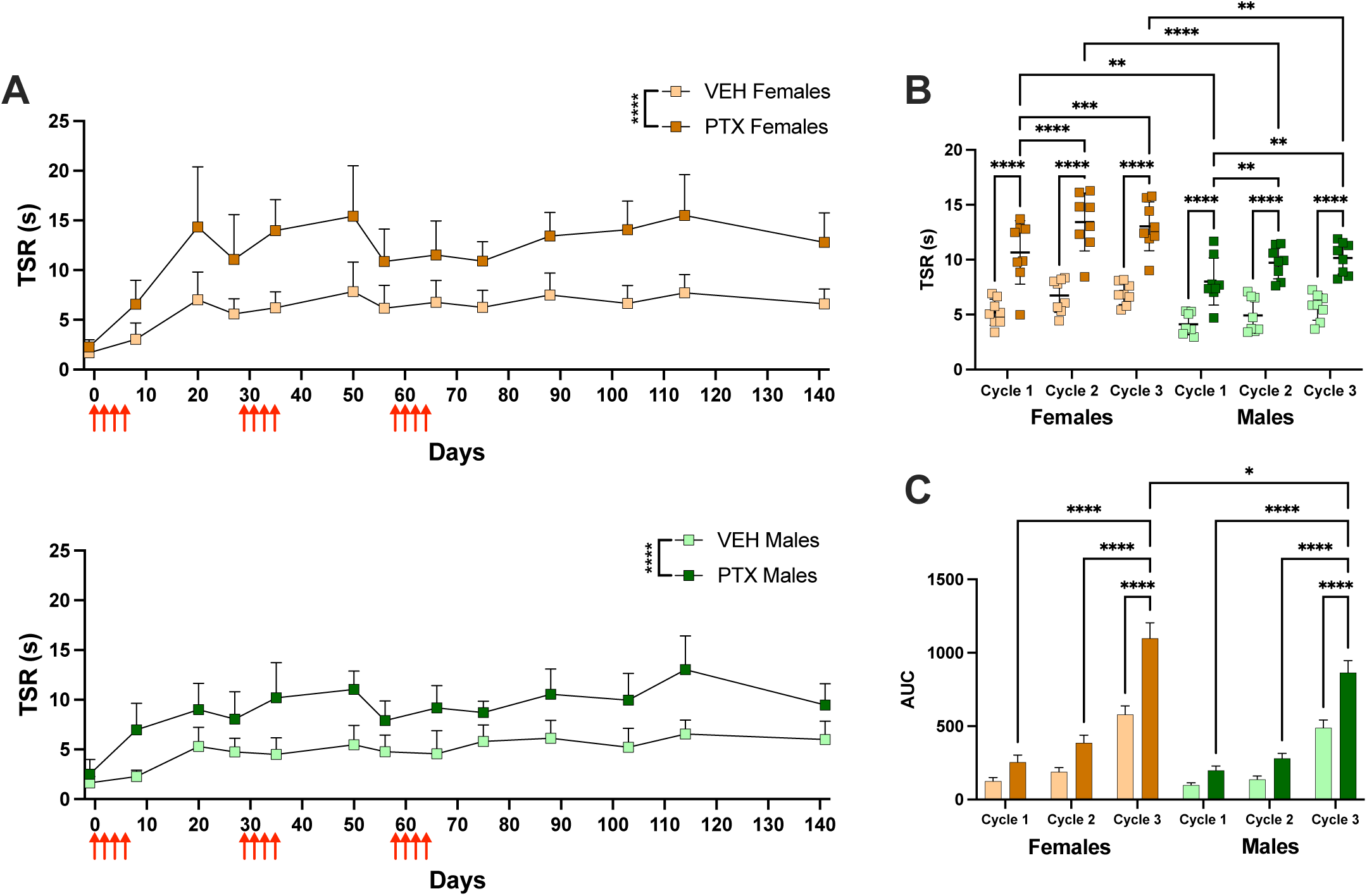
Increased duration of cold hypersensitivity after multiple PTX cycles. **A)** Time spent reacting (seconds) to acetone stimulation throughout administration of 3 PTX cycles (red arrows) in female (top) and male (bottom) mice. **B)** Average time spent reacting (seconds) throughout each cycle of PTX administration in female and male mice. **C)** Area under the curve (AUC) of cold sensitivity to acetone following the administration of each PTX cycle in female and male mice. (n=8/group). Data is represented as mean±SD. * p < 0.05, ** p < 0.01, *** p<0.001, and **** p < 0.0001.

### 3.5 Repeated PTX cycles do not inhibit proprioception or motor function

We used the rotarod to examine changes in proprioception and motor function due to repeated PTX cycles. We found no differences in the latency to fall from the rotarod, or the RPM at fall, between PTX and vehicle groups for male or female mice (Fig. 6A, Supp Fig. 3). However, both male [*F*(5.617, 76.53) = 24.16, p<0.001] and female [*F*(4.842, 65.67) = 30.01, p<0.0001] mice showed an increase in ability to stay on the rotarod as the test was repeated over time (Fig. 6A).

**Figure 6.**
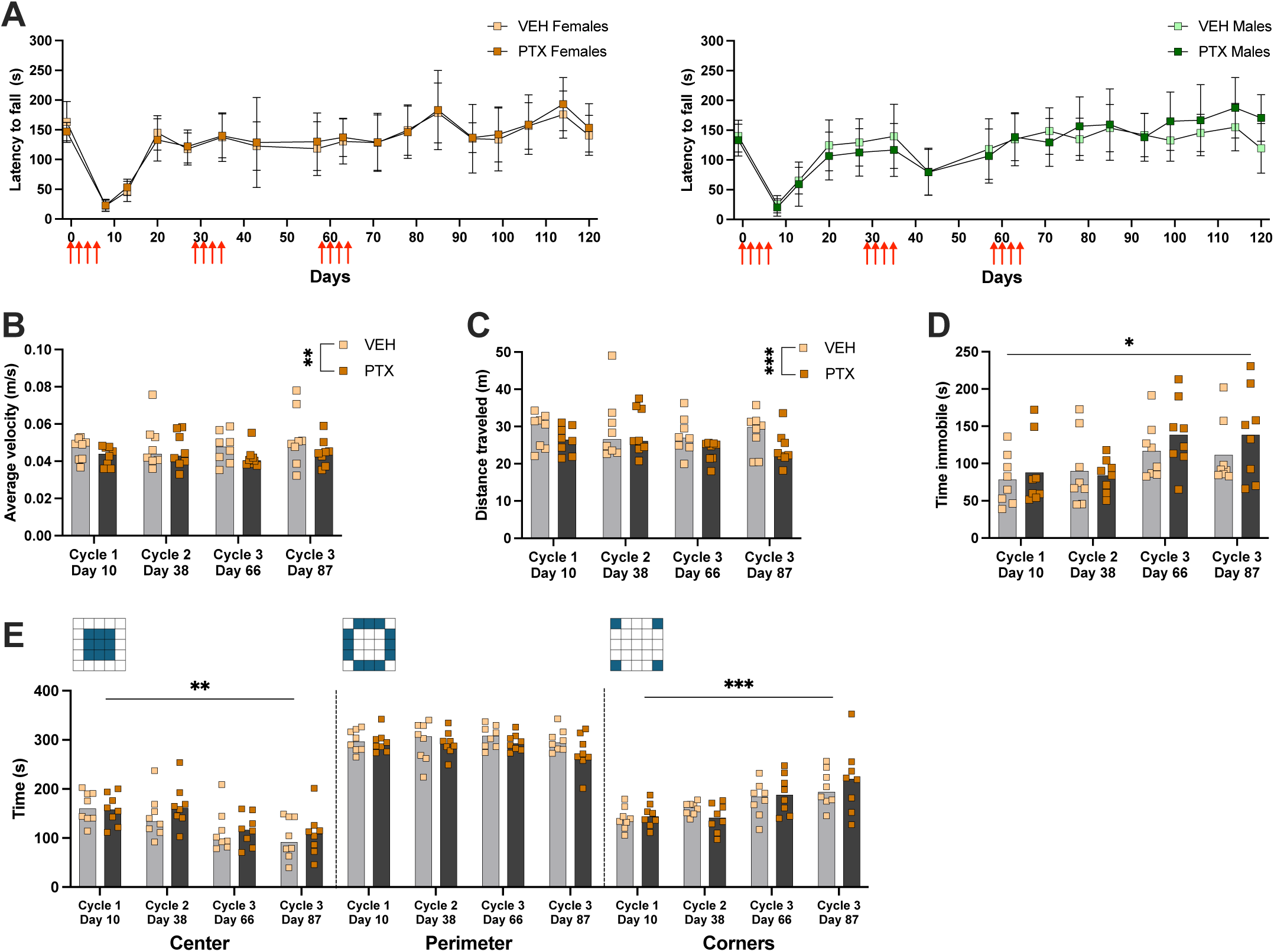
Proprioception and motor function after multiple PTX cycles. **A)** Latency (seconds) to fall from rotarod throughout administration of 3 PTX cycles (red arrows) in female (left) and male (right) mice. **B)** Average velocity (meters/second), **C)** distance traveled (meters), **D)** time spent immobile (seconds), and **E)** time spent in designated areas (seconds) in open field apparatus following each PTX cycle in female mice. Mice were only tested once in the open field apparatus. (n=8/group). Data is represented as mean±SD. * p < 0.05, ** p < 0.01, and *** p<0.001.

To examine voluntary mobility and anxiety-like behaviors we tested female mice in the open field test during the peak of mechanical hypersensitivity between days 7-10 following the first dose of each cycle of PTX, as well as 30 days following the first dose of the third cycle of PTX. Importantly, each mouse was only tested once. PTX significantly decreased the average velocity [*F*(1, 14) = 10.57, p=0.0058] and distance traveled [*F*(1, 14) = 17.93, p=0.0008] (Figs 6B, C), indicating that PTX induced a decrease in voluntary mobility. We found a significant effect of time on the time spent immobile in the open field [*F*(2.419, 33.86) = 4.030, p=0.0208] (Fig. 6D). We also found that over time, mice decreased time spent in the center [*F*(2.376, 33.27) = 7.745, p=0.001] and increased time spent in the corner areas [*F*(2.144, 30.02) = 10.92, p=0.0002] of the open field (Fig. 6E), suggesting a decrease in exploratory behaviors. There was no change in time spent on the perimeter of the open field (Fig. 6E). To assess potential sex differences, we tested male mice in the open field 30 days following the third cycle of PTX. We found no differences in time spent immobile, average velocity, time spent in specific areas of the open field, or distance traveled between the PTX and vehicle groups (Supp Fig 4).

## 4. Discussion

In this study, we introduce a translational rodent CIPN model that uses three consecutive PTX cycles to generate a clinically relevant phenotype of persistent mechanical and cold hypersensitivity. Our comprehensive, longitudinal behavioral assessments identify the effects of repeated PTX cycles on PIPN development, resolution, and chronification. Through measurements of mechanical sensitivity, we demonstrated that repeated cycles of PTX significantly increases the duration of mechanical hypersensitivity. We showed that PTX induces significant cold hypersensitivity that does not develop or resolve along the same trajectory as mechanical hypersensitivity. Further, we found that repeated cycles of PTX administration do not impair proprioception or motor function but reduce voluntary mobility in female mice in the open field apparatus.

PTX is known to cause serious adverse effects (e.g., neutropenia, anemia, nausea and vomiting, loss of appetite, weight loss, myocardial toxicity) that worsen with repeated cycles of administration [19,21,26,31]. Current rodent models of PIPN administer a cumulative PTX dose of 8-32 mg/kg for a single cycle and do not demonstrate many of these adverse effects [17,37]. As we administered three cycles of PTX with a cumulative dose of 48 mg/kg, we sought to monitor the mouse’s overall wellbeing. Slowing of weight gain in models of CIPN has not been consistently reported [37,39]. In our study, PTX did not alter food consumption for male or female mice but did affect weight gain in males, indicating that PTX has some physiological effects in male mice.

Hematologic abnormalities (e.g., anemia, leukopenia, and thrombocytopenia) commonly affect cancer patients receiving PTX [23,26] and are reported in rodent PTX models [28]. In the study by Park et al, a dose of 8.5 mg/kg PTX was administered daily for five days and induced anemia and leukopenia compared to a PBS control in BALB/c mice. In our study, both PTX and vehicle groups showed RBC and HGB slightly lower than the normal range, indicating anemia, WBC lower than the normal range, indicating leukopenia, and PLT lower than the normal range, indicating thrombocytopenia. PTX did not alter complete blood count results compared to the vehicle control, possibly due to changes occurring from the Cremophor EL vehicle. Our schedule of repeated PTX cycles did not impact hematologic measurements over time. Consistent with prior findings, PTX did not impact normal rodent burrowing behavior [39]. Overall, we demonstrate that repeated cycles of PTX do not have large negative impacts on male or female mice overall health and can therefore be used to examine PIPN in rodents without confounding physiological effects.

Mechanical hyperalgesia is one of the hallmark features of CIPN which limits the use of chemotherapy in cancer treatment and decreases the quality of life for cancer survivors [9,25]. Therefore, preclinical models which closely mirror these clinical features are essential in understanding CIPN pathogenesis and developing effective treatment options. Current rodent models of PIPN measure mechanical sensitivity following a single cycle of PTX where hypersensitivity develops within 14 days and resolve after 28 days [13,17,37]. We are aware of a single report utilizing more than one PTX cycle in rodents. Toma and colleagues administered a second cycle of PTX one week after the initial cycle and demonstrated a further decrease in mechanical threshold in the second cycle compared with the first, as well as an extension of mechanical hypersensitivity from 28 days to 42 days [37]. In the present study, PTX produced mechanical hypersensitivity in male and female mice that developed and resolved following a single cycle of PTX consistent with previous findings [13,17,37]. The intensity and duration of hypersensitivity following a second PTX cycle was similar to that of the first cycle. While these findings differ from those reported by Toma et al, the difference in timing of the second PTX cycle may explain this inconsistency. Notably, we show a 2-fold increase in the duration of mechanical hypersensitivity following the third PTX cycle in both males and females. This previously unreported phenotype of mechanical hypersensitivity mirrors the chronification of CIPN seen clinically. The underlying mechanisms of this long-lasting hypersensitivity are not known and should be investigated in future studies.

Cold allodynia is another major symptom of CIPN that contributes to patient discontinuation of cancer treatment [31,36] and therefore must be considered when examining CIPN in rodents. While cold hypersensitivity develops in preclinical CIPN models 7-10 days after the first dose [37], its resolution is not consistently reported. Warnke et al showed cold hypersensitivity following a single cycle of 3 mg/kg oxaliplatin lasted for 10 weeks [39]. A cycle of 1 mg/kg PTX induced cold hypersensitivity that persisted for >30 days in male ICR mice [8] while a cycle of 2 or 4 mg/kg PTX induced cold hypersensitivity that resolved by day 22 in male C57BL/6J mice [37]. Our findings are consistent with the reported development of CIPN in rodents as cold hypersensitivity developed within 7-10 days following the first PTX cycle [8,37,39]. Variability in reports of cold hypersensitivity resolution may be due to differences in the use and scoring of behavioral assays [3,12]. We do not have knowledge of prior studies that measure cold sensitivity during repeated chemotherapeutic cycles. Consistent with the cumulative effects of PTX observed clinically [36], our model produces persistent cold hypersensitivity that increases with subsequent cycles. Unlike responses to mechanical stimuli, females had increased intensity of cold hypersensitivity compared to males due to PTX, demonstrating sensory specific sex differences in PIPN. Further, cold hypersensitivity persisted even 12 weeks after the third cycle of PTX was given, which is considerably longer than the 8-week duration of mechanical hypersensitivity after the third cycle. These differences in mechanical and cold hypersensitivity development and resolution in CIPN suggest that both modalities should be considered in preclinical rodent models to properly characterize CIPN pathogenesis and evaluate potential therapies. Future studies should examine the mechanisms of sensory-specific dysregulation occurring following repeated chemotherapy cycles. Collectively, these findings demonstrate that repeated PTX cycles in rodents induces long-lasting hypersensitivities that mirror the clinical persistence of mechanical hyperalgesia and cold allodynia in CIPN.

Patients with CIPN may also experience impaired mobility (e.g., balance and postural control, upper limb function, gait) [38]. While CIPN does not impact wheel running or rotarod performance in rodents [3,39], gait disorders have been reported [22]. Mice that received viscristine or bortezomib had a shorter distance traveled in the open field test [3], while those given doxorubicin spent less time in the center of the open field [6]. Consistent with these findings, we found that PTX reduced velocity and distance traveled in the open field, but did not impair rotarod performance in our model [3,6,39]. Our study design enabled us to assess changes in open field behavior throughout repeated PTX cycles. There was a decrease in time spent in the center with subsequent cycles in both groups. All mice receive Cremophor EL as a component of the vehicle; Cremophor EL is a known neurotoxin and may contribute to behavioral changes [14]. These findings may also indicate age-related changes in behavior as we tested mice from 8-20 weeks of age in the open field. Further studies are needed to examine the effects of Cremophor EL and age on these behaviors are needed to properly assess how PTX alters exploratory and anxiety-like behaviors. Overall, this model of repeated PTX cycles induces a phenotype that does not impair proprioception or motor function but does impact voluntary movement, mimicking cases where CIPN interferes with patient mobility and activities of daily life [36,38].

## Conclusion

We show that repeated cycles of PTX, which mimics the clinical dosing regimen, induces a phenotype similar to the clinical presentation of CIPN with long-lasting cold and mechanical hypersensitivity in rodents without impaired proprioception and motor function. We found that the cumulative effect of PTX induces persistent physiological changes despite mechanical hypersensitivity resolution between cycles. The differences in the development and persistence of cold and mechanical hypersensitivity in our model illustrates the importance of an independent evaluation of these behaviors and their underlying mechanisms in CIPN. We anticipate that the use of this model will improve our understanding of CIPN chronification and will facilitate the development of translational therapies for CIPN. Effective pain management will enable patients to complete their cancer treatment, decrease health care expenditure, decrease mortality, and improve the physical and mental health and quality of life of cancer patients and survivors.

## 5. Author contributions

**LRO**: Conceptualization, Methodology, Investigation, Writing – Original Draft. **MGB, KAR, DMR**: Investigation, Writing – Review and Editing. **EAD:** Writing – Review and Editing**. KES**: Conceptualization, Investigation, Resources, Writing – Review and Editing.

## 6. Acknowledgements

This study was funded by the National Institutes of Health (P20GM121293), the Arkansas Children’s Research Institute, the University of Arkansas for Medical Science Winthrop P. Rockefeller Cancer Institute, and a Seeds of Science research grant. The funders had no role in the study design, data collection and analysis, decision to publish, or preparation of the manuscript. The authors have no conflicts of interest to declare.

