## Supplemental information for "A novel model of paclitaxel-induced peripheral neuropathy produces a clinically relevant phenotype in mice"

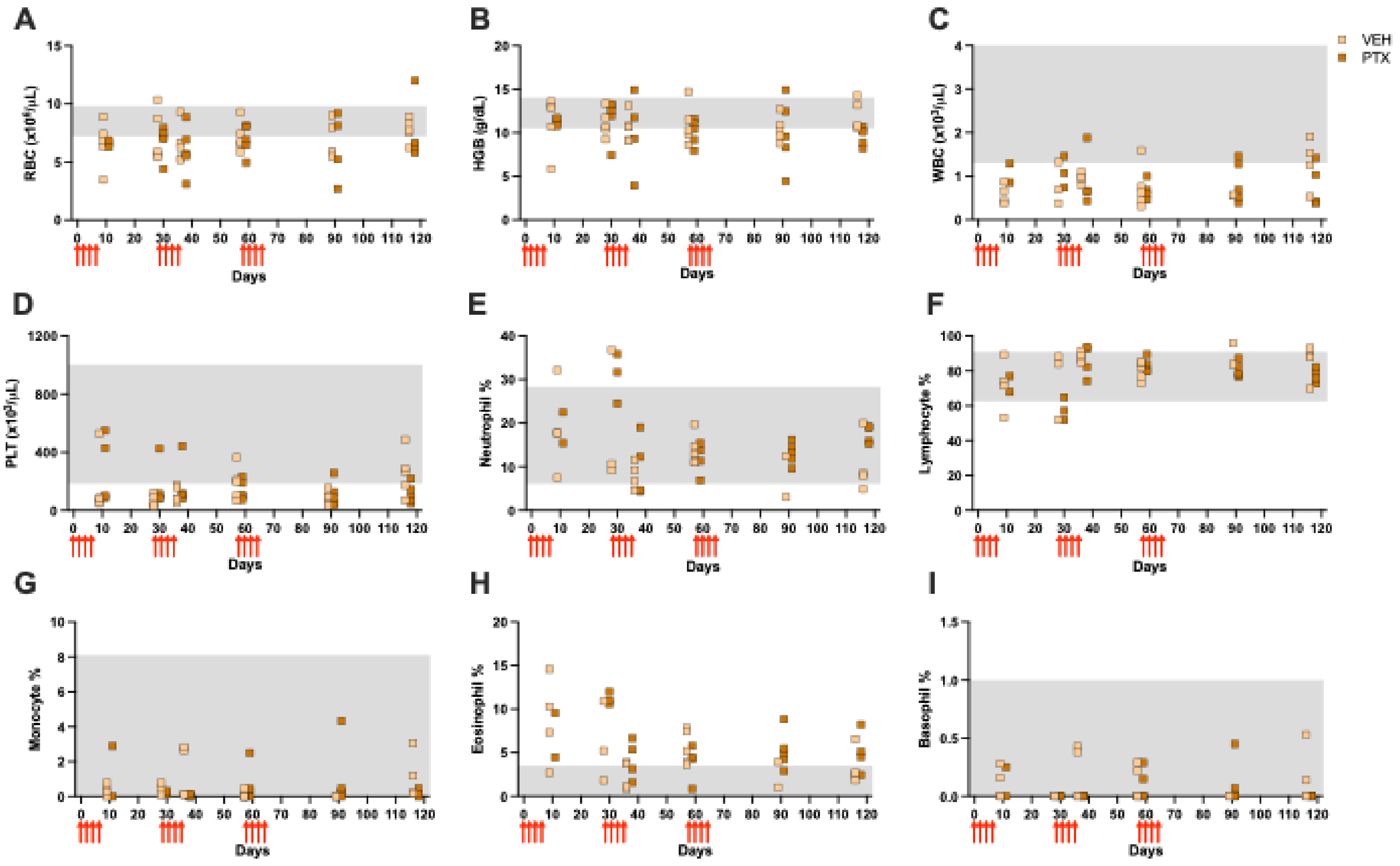


**Supplementary Figure 1. Complete blood count of female mice throughout study. A)** red blood cell count ($x10^6/\mu L$), **B)** hemoglobin (g/dL), **C)** white blood cell count ($x10^3/\mu L$), **D)** platelet count ($x10^3/\mu L$), **E)** neutrophil %, **F)** lymphocyte %, **G)** monocyte %, **H)** eosinophil%, and **I)** basophil % throughout administration of three cycles of PTX (red arrows). Normal range of values indicated by shaded areas. (n=5/group).

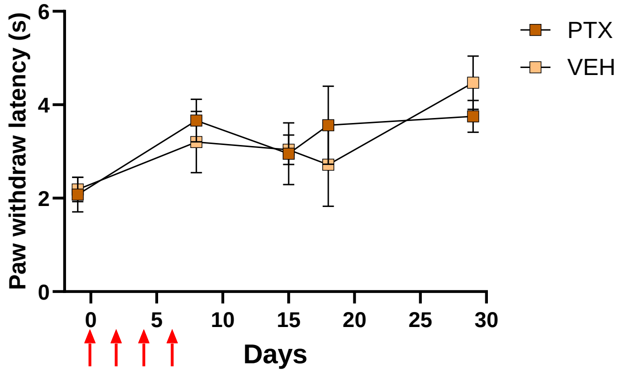


**Supplementary Figure 2. Thermal sensitivity after a single PTX cycle.** Paw withdrawal latency (seconds) to thermal stimulation following the administration of one PTX cycle (red arrows) in female mice. (PTX, *n=5*; VEH, *n=4*). Data is represented as mean±SD.

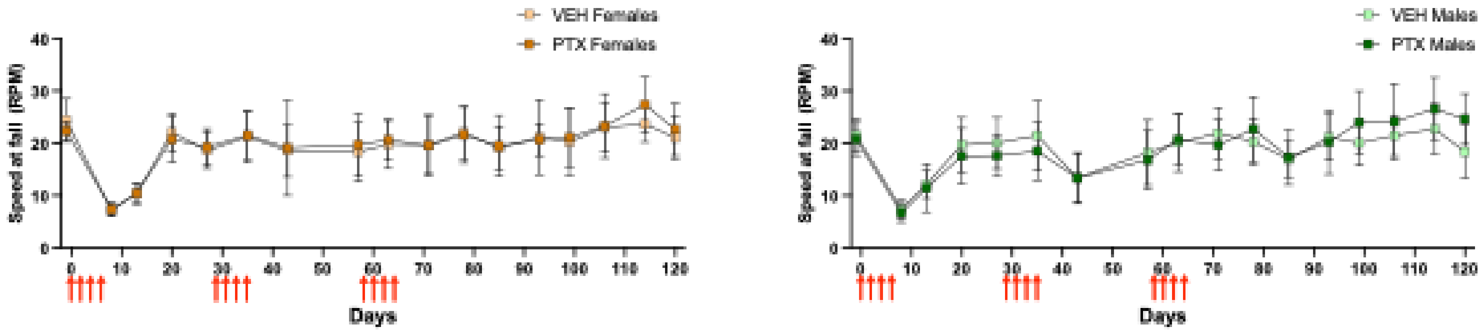


**Supplementary Figure 3. Proprioception after multiple PTX cycles.** Speed of rotarod spinning (rpm) at fall from rotarod throughout administration of 3 PTX cycles (red arrows) in female (left) and male (right) mice. (*n=8* per group) Data is represented as mean±SD.

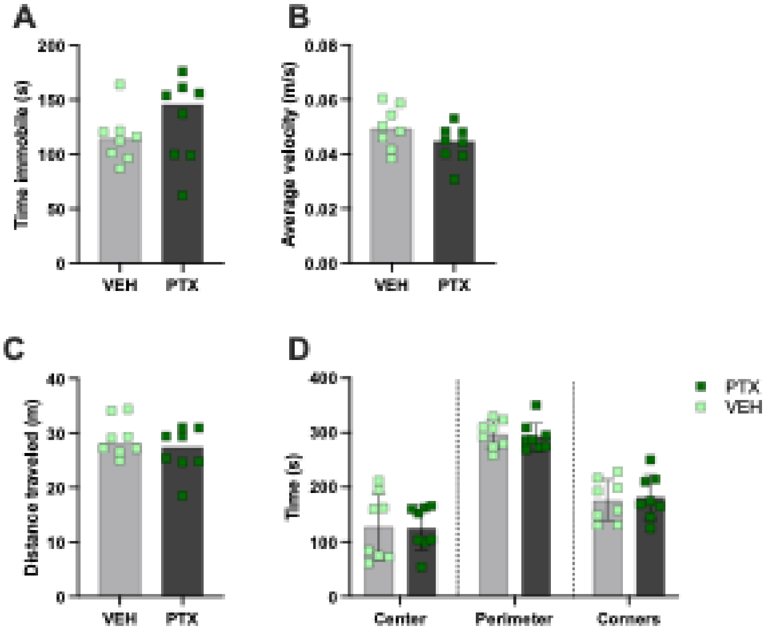


**Supplementary Figure 4. Motor function after multiple PTX cycles. A)** Time spent immobile (seconds), **B)** average velocity (meters/second), **C)** distance traveled (meters) and **D)** time spent in designated areas (seconds) in the open field following administration of the third cycle of PTX or vehicle in male mice (day 87). (*n=8* per group, mice were only tested once). Data is represented as mean±SD.
